# A novel expression system enabling scalable production of glycosylated flavonoids in *Escherichia coli* W using a plant-derived toxic gene

**DOI:** 10.1101/2025.09.09.674713

**Authors:** Darwin Carranza-Saavedra, Jesús Torres-Bacete, Juan Nogales

## Abstract

Glycosylated flavonoids are plant-derived compounds of significant interest due to their enhanced solubility, stability, and bioavailability, offering therapeutic potential across pharmaceutical, nutraceutical, and cosmetic sectors. However, their complex biosynthesis in plants hinders scalable production. In this study, we present an innovative microbial platform based on a phosphate-responsive *Pliar*53 promoter system in non-model chassis *Escherichia coli* W to enable efficient, regulated heterologous expression of the *Sbai*C7OGT gene from *Scutellaria baicalensis*. This platform circumvents common limitations associated with conventional inducible systems that rely on costly or toxic chemical inducers. The engineered *E. coli* SBG2413 strain demonstrated high titers of naringenin-7-O-glucoside (prunin) and exhibited broad substrate compatibility for the biosynthesis of other flavonoid glycosides. Our findings establish a cost-effective, scalable solution for industrial production of glycosylated flavonoids, with potential applicability to co-culture systems and microbial consortia.

**Highlights:**

- A phosphate-response gene expression system (*Pliar53*) enables stable expression of a toxic flavonoid glycosyltransferase.
- Translational optimization using the bicistronic BCD2 RBS and high-copy plasmids boosts prunin production from naringenin.
- *E. coli* W SBG2413strain achieves > 9 g/L prunin in fed-batch bioreactor with 70% conversion.
- The platform developed allows efficient glycosylation of structurally diverse flavonoids including flavones, flavanones, and isoflavones.

## 1. Introduction

Flavonoids, a major class of polyphenolic secondary metabolites, are widely acknowledged for their broad spectrum of biological activities, including antioxidant, anti-inflammatory, and anticancer effects (Ribeiro et al., 2023). However, their inherently low aqueous solubility presents significant challenges for their application as industrial bioproducts. To overcome this limitation, the glycosylation of flavonoids can significantly improve their physicochemical properties such as solubility, stability, and bioavailability, which significantly increase their suitability for industry use (Mrudulakumari Vasudevan & Lee, 2020). Despite these advantages, the large-scale production of glycosylated flavonoids remains limited by the complexity of their biosynthesis in plants and the inefficiencies of traditional heterologous expression systems in microbial hosts.

Flavonoid glycosylation has been pursued through several strategies, including *in vitro* enzymatic conversion using purified UDP-glycosyltransferases (UGT), whole-cell biocatalysis, and microbial metabolic engineering (Feng et al., 2020; Xiao et al., 2014). A central requirement for all these approaches is the availability of UDP-glucose (UDPG), the universal sugar donor in flavonoid glycosylation reactions (J. Li et al., 2025; Matera et al., 2024). Consequently, ensuring a sustainable and cost-effective intracellular supply of UDPG is critical for the viability of industrial-scale bioprocesses (Wang et al., 2022).

In parallel, the efficiency of these biotechnological strategies also depends on the successful heterologous expression of plant-derived enzymes in microbial hosts such as *Escherichia coli* (Mrudulakumari Vasudevan & Lee, 2020). Among these, *E. coli* W, a less commonly used but metabolically versatile strain, has gained attention for its robustness and capacity to accommodate non-native biosynthetic pathways (Carranza-Saavedra et al., 2021, 2023; Erian et al., 2018; Felpeto-Santero et al., 2015). Nevertheless, expressing plant enzymes in *E. coli* often poses challenges due to the lack of compatible folding environments, post-translational modifications, or essential cofactors.

In response to these bottlenecks, various strategies have been employed, including codon optimization, chaperone co-expression, fusion protein engineering, and inducible promoter systems (Watts et al., 2021). While promising, these methods often yield inconsistent results and require case-by-case optimization. A key element in recombinant protein production is the regulation of gene transcription (Watts et al., 2021). These limitations underscore the urgent need for advanced expression systems that provide tight, tunable control over plant gene expression in microbial hosts, offering both scalability and robustness for industrial deployment (Lee et al., 2024). Among regulatory systems, the *XylS*/*Pm* system, inducible by m-toluic acid, and the *RhaR/Prha* system, activated by L-rhamnose, have been widely applied in metabolic engineering and synthetic biology because these regulons are notable for their tight transcriptional control and high expression levels (Kim et al., 2025; Matera et al., 2025). However, both systems rely on costly chemical inducers and are associated with potential toxicity and poor scalability. For example, m-toluic acid derivatives such as 3-methylbenzoate (3 MB) are toxic and can interfere with enzymatic reactions (Mohanapriya et al., 2016; Naz et al., 2022), while the high cost of L-rhamnose limits its use in large-scale processes (Rong et al., 2024). These limitations underline the need for more economical and scalable alternatives.

To address these limitations, the synthetic phosphate (Pi)-depletion promoter system, known as *Pliar*, has been proposed as a cost-effective, scalable, and environmentally friendly alternative to conventional chemically inducible systems (Torres-Bacete et al., 2021). Unlike classical expression systems such as *LacI/Ptrc* the *XylS*/*Pm* and *RhaR*/*Prha* systems, which require expensive and potentially toxic inducers, the *Pliar* system enables automatic, tightly regulated gene expression in response to phosphate availability in the medium (Torres-Bacete et al., 2021). This system is inspired by the bacterial phosphate (PHO) starvation response, in which low Pi concentrations activate the PHO regulon. Under these conditions, the transcriptional regulator *Pho*B becomes phosphorylated and initiates the expression of phosphate-responsive genes (Fig. 1A and 1B). Synthetic promoters engineered from native PHO-regulated promoters (such as *pst*S, *pho*A, or *pho*B) contain conserved PHO boxes and drive transcription only when phosphate concentrations fall below a specific threshold (Yu et al., 2023). Thus, the Pi levels can be precisely controlled through medium formulation, the *Pliar* system offers a low-cost, non-toxic induction mechanism, eliminating the need for compounds such as 3 MB or L-rhamnose. As a result, *Pliar* represents a versatile and industrially relevant expression system, ideally suited for the biosynthesis of valuable biochemicals like glycosylated flavonoids, where fine-tuned gene expression is essential for productivity and host stability.

**Fig. 1.**
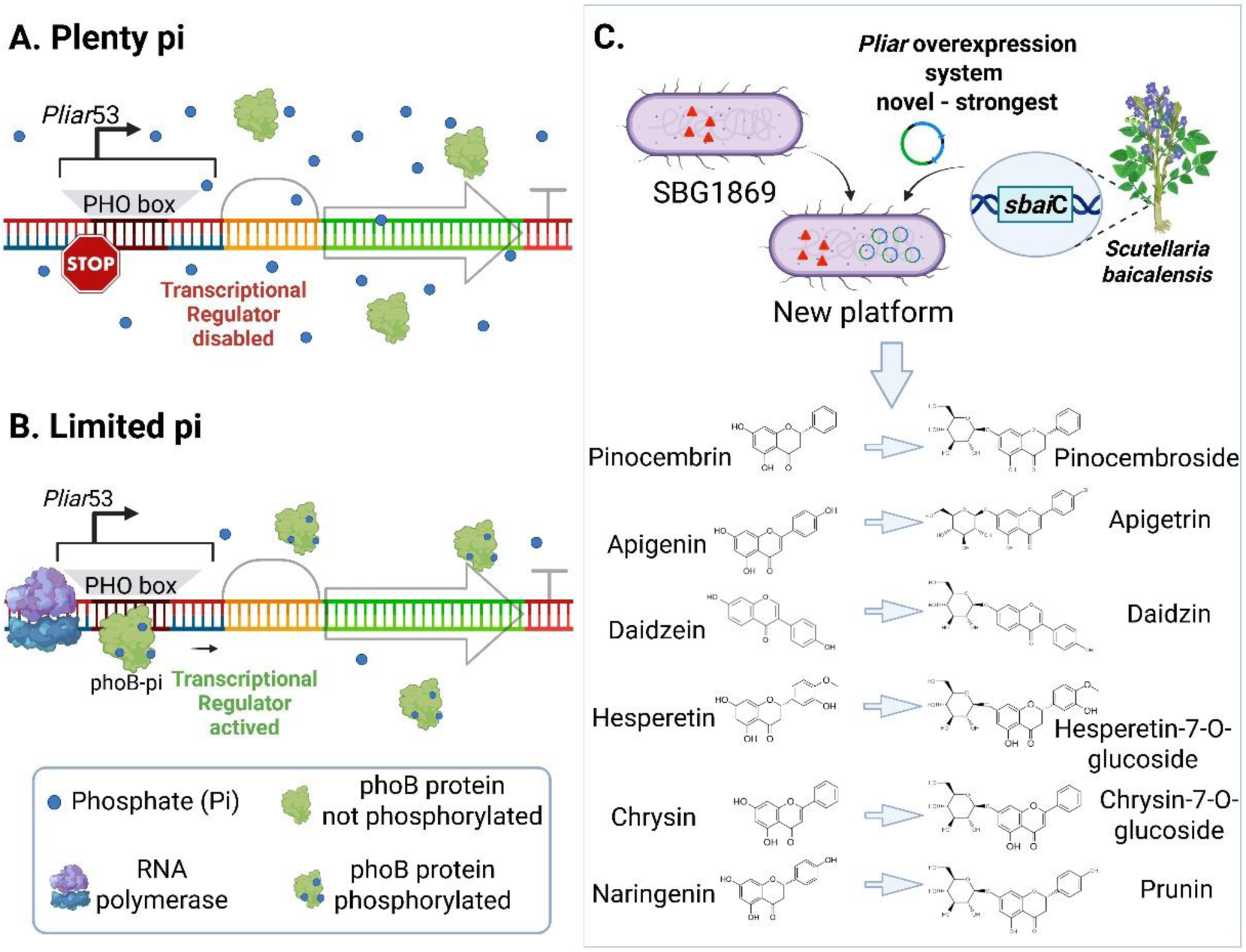
Functional scheme of the *Pliar* overexpression system used in this study. **A.** Under high-phosphate (pi) conditions, transcription from the *Pliar53* system is repressed. **B.** Under phosphate-limiting conditions, the *Pliar53* system is activated, enabling gene expression. **C.** Overview of the conceptual framework and overall strategy employed in this study.

In this context, the *Pliar* system represents a promising strategy for the tightly regulated expression of potentially toxic genes involved in flavonoid glycosylation. However, its potential for enabling the biosynthesis of complex biomolecules such as flavonoids remains largely unexplored. Moreover, evaluating its true effectiveness requires a host platform capable of reliable and high level UDPG production. In previous work, we successfully engineered *E. coli* W to metabolize sucrose (a low-cost and abundant carbon source) for efficient UDPG biosynthesis and subsequent glycosylation reactions (Carranza-Saavedra et al., 2025). Nevertheless, the glycosylation potential of this strain had not yet been systematically explored. To explore this capability, we investigated the glycosylation of naringenin to produce prunin using the *SbaiC*7OGT gene from *Scutellaria baicalensis* (Matera et al., 2024), a glycosyltransferase well characterized for its specific activity in 7-O-glucosylation of diverse flavonoid substrates (Hirotani et al., 2000). This enzyme’s regioselectivity offers the advantage of producing a single glycosylated product per substrate, thereby simplifying downstream purification in bioprocess applications, although it has been reported to be toxic for host cells. To address this, we employed the *Pliar* system, which enables phosphate-responsive and leaky expression control, to minimize potential toxic effects. Using this strategy, we achieved the highest reported titer of prunin through bioprocess scale-up.

Finally, to demonstrate the robustness and versatility of our engineered platform, we expanded its application to include a range of structurally distinct flavonoids (Fig. 1C). The platform successfully glycosylated pinocembrin, apigenin, daidzein, chrysin, and hesperetin, confirming its broad substrate scope and industrial potential for the scalable production of flavonoid glycosides.

## 2. Material and methods

### 2.1. Strains, plasmids and DNA parts

The strains, plasmids, and DNA parts utilized in this study are detailed in Table 1.

**Table 1.** Strains and plasmids used.

| Strain | Genotype | Source |
| --- | --- | --- |
| *E. coli* DH5α | fhuA2 Δ(*argF-lacZ*)U169 *phoA* *glnV44* Φ80 Δ(*lacZ*)M15 *gyrA96* *recA1* *relA1* *endA1* *thi-1* *hsdR17* | New England BioLabs (NEB) |
| *E. coli* SBG1869 | *E. coli* SBG1103, ALE sucrose, Δ*xylA* Δ*zwf* Δ*pgi* Δ*glgC* | (Carranza-Saavedra et al., 2025) |
| *E. coli* SBG1882 | *E. coli* SBG1869, pOSR | This study |
| *E. coli* SBG1883 | *E. coli* SBG1869, pYSR | This study |
| *E. coli* SBG1888 | *E. coli* SBG1869, pUDG | This study |
| *E. coli* SBG2283 | *E. coli* SBG1869, pNSR | This study |
| *E. coli* SBG2303 | *E. coli* SBG1888, pSSR | This study |
| *E. coli* SBG2339 | *E. coli* SBG1888, pPSS | This study |
| *E. coli* SBG2338 | *E. coli* SBG1888, pPBSr | This study |
| *E. coli* SBG2412 | *E. coli* SBG1888, pPBSb | This study |
| *E. coli* SBG2413 | *E. coli* SBG1888, pPBSu | This work |
| Plasmid | Description | Source |
| pUDG | Gm<sup>r</sup>, pUC ori, CDS: *XylS/Pm-glK-pgm-galU*, RBS-ST, BBa_B1006 terminator and linkers | This study |
| pOSR | Gm<sup>r</sup>, RK2 ori, CDS: *XylS/Pm-glK-pgm-galU-sbaiC*, RBS-ST, BBa_B0014 terminator and linkers | This study |
| pYSR | Gm<sup>r</sup>, RK2 ori, CDS: *RhaR/Prha-glK-pgm-galU-sbaiC*, RBS-ST, BBa_B0014 terminator and linkers | This study |
| pSSR | Km<sup>r</sup>, RK2 ori, CDS: *Pm-sbaiC*, RBS-ST, BBa_B1006 terminator and linkers | This study |
| pRSR | Km<sup>r</sup>, RK2 ori, CDS: *RhaR/Prha-sbaiC*, RBS-ST, BBa_B1006 terminator and linkers | This study |
| pPSS | Km<sup>r</sup>, RK2 ori, CDS: *Pliar53-sbaiC*, RBS-ST, BBa_B1006 terminator and linkers | This study |
| pPBSr | Km<sup>r</sup>, RK2 ori, CDS: *Pliar53-sbaiC*, RBS-BCD2, BBa_B1006 terminator and linkers | This study |
| pPBSb | Km<sup>r</sup>, pBBR ori, CDS: *Pliar53-sbaiC*, RBS-BCD2, BBa_B1006 terminator and linkers | This study |
| pPBSu | Km<sup>r</sup>, pUC ori, CDS: *Pliar53-sbaiC*, RBS-BCD2, BBa_B1006 terminator and linkers | This study |

### 2.2. DNA manipulation

Plasmids and DNA fragments were purified using commercial kits supplied by NZYtech (Lisbon, Portugal). Oligonucleotides were supplied by Merck (Darmstadt, Germany). Restriction enzymes used were supplied by New England Biolabs (Ipswich, USA), except BpiI, which was sourced from Thermoscientific (Waltham, USA).

The codifying sequences (CDSs) for the *glk*, *pgm*, *gal*U genes (*E. coli* W), and *Sbai*C7OGT (*S. baicalensis*) genes were domesticated manually by removing BsaI and BpiI restriction sites and then were constructed by Explora Biotech Srl company (Italy). All genetic constructions were confirmed by sequencing (Eurofins Genomic, Germany). We designed the synthetic operons for production of UDPG and glycoside flavonoids (Table 1) by Modular Cloning (MoClo), using our lab’s Golden-Standard platform (Blázquez et al., 2023).

### 2.3. Reagents, culture media and conditions

Naringenin, hesperetin and chrysin were purchased from Sigma-Aldrich (Darmstadt, Germany). Prunin (naringenin-7-O-glucoside), daidzein, daidzin (Daidzein-7-O-glucoside), apigenin, apigetrin (apigenin-7-O-glucoside), hesperetin-7-O-glucoside, pinocembrin, pinocembroside (pinocembrin-7-O-glucoside) and Chrysin-7-O-Glucoside were acquired from Extransynthesis (Lyon, France). Other reactive agents used were analytical grade. All the flavonoids were resuspended with dimethyl sulfoxide (DMSO). *E. coli* strains were grown in Luria-Bertani (LB) medium, preinocula and production mediums: M9 medium (pH 7.2) (Landor et al., 2023) or M3 medium supplemented with MOPS buffer at pH 7.2, 3 g/L yeast extract, and devoid of phosphate (Neidhardt et al., 1974). All production mediums were supplemented with 0.034 mM EDTA, 3 mM flavonoid, and 1.5 mM 3-Methylbenzoate (3MB) or 1.0 mM rhamnose inducer, gentamicin (20 µg/mL) and/or kanamycin (50 µg/mL) when was necessary. All culture media and work solutions were sterilized at 121°C for 15 min and their initial pH was adjusted to 7.0-7.2 or carefully filtered using a 0.2 µm cellulose nitrate filter before using it.

### 2.4. Production in shake flasks and bioreactor

The production of flavonoid glycosides was conducted in both shake flasks and bioreactor systems. For flask experiments, strains were first cultured in 10 mL of LB medium (seed culture) in 50 mL Erlenmeyer flasks and incubated at 37 °C and 250 rpm in a MaxQ 4000 orbital shaker (Thermo Scientific, United States) for 6-8 hours. Subsequently, 2 mL of seed culture were transferred to 100 mL of M9 medium containing 15 g/L sucrose in 500 mL shake flasks and incubated at 37 °C and 250 rpm for 15-18 hours (pre-inoculum stage). Cells were then harvested by centrifugation, resuspended in 0.85% (w/v) NaCl, and inoculated into 100 mL of either M9 or M3 production medium (with 10 g/L sucrose) in 500 mL flasks at an optical density at 600 nm (OD600) of 0.5 - 2.0. Cultures were incubated at 37 °C and 250 rpm for the production phase.

For bench scale-up experiments, glycosylation was performed in a custom-built 5 L bioreactor (Bioprocess Technology S.L., Spain) with a working volume of 2 L, operated with a low-cost production medium. A total of 400 mL of pre-inoculum grown in M9 medium was harvested by centrifugation at 5,200 × g for 10 min, resuspended in 0.85% (w/v) NaCl, and transferred into the bioreactor containing 1.5 L of medium formulated without phosphate and MOPS buffer. The base medium per liter of distilled water contained: 10.00 g sucrose, 0.50 g NaCl, 6.00 g NH_4_Cl, 5.00 g yeast extract and trace metals prepared in 1 N HCl (2.78 mg FeSO_4_·7H_2_O, 1.98 mg MnC_2_·4H_2_O, 2.38 mg CoCl_2_·6H_2_O, 1.47 mg CaCl_2_·2H_2_O, 0.25 mg CuSO_4_·5H_2_O, 0.29 mg ZnSO_4_·7H_2_O). A Silicone Antifoam (30% in H_2_O) was added at 600 µL per batch. The bioprocess was carried out at 37 °C with automatic pH control set at 7.2, adjusted using 2 M NaOH. Dissolved oxygen was maintained at approximately 20% saturation by modulating the agitation speed between 300 and 800 rpm with aeration of 1 vvm, as described in Carranza-Saavedra et al. (2021, 2025).

Fed-batch operation was carried out using concentrated feedstocks: 500 g/L sucrose solution was added intermittently by pulse to maintain a final sucrose concentration of 10–15 g/L; 200 g/L yeast extract was supplemented to maintain a final concentration of 3 g/L. Naringenin (190 g/L in DMSO) was introduced progressively into the bioreactor culture through a stainless steel capillary tube (Carranza-Saavedra et al., 2025) using a MINIPULS 3 peristaltic pump (Gilson, Spain), targeting a final concentration of 0.8 g/L. Real-time monitoring of cell growth and sucrose consumption was employed to optimize the timing of naringenin and sucrose addition and mitigate potential cytotoxic effects, thereby enhancing overall process efficiency.

### 2.5. Analytical methods

Bacterial growth was monitored by measuring OD600 using a Ubet-30 spectrophotometer (Jasco, Japan). Cell dry weight per liter (g CDW/L) was determined using the formula: 1 g CDW/L = OD600 × 0.452 (Erian et al., 2018).

Flavonoid quantification was carried out based on the method previously described by Carranza-Saavedra et al. (2025), with minor modifications. Separation was performed using high-performance liquid chromatography (HPLC) with a diode array detector (DAD). The mobile phase consisted of methanol (solution A) and water containing 0.1% formic acid (solution B), with a flow rate of 1.0 mL/min. The following gradient program was applied: starting at 40% A, increasing to 43% A over 5 min, then to 70% A over the next 11 min, and finally reaching 95% A for 1 min. Detection was performed using a DAD detector at specific wavelengths: 250 nm for daidzein and daidzin, 283 nm for prunin, 284 nm for hesperetin-7-O-glucoside and pinocembroside, 288 nm for hesperetin and pinocembrin, 290 nm for naringenin, 304 nm for chrysin-7-O-glucoside, 312 nm for chrysin, and 337 nm for apigenin and apigetrin. Analysis of sucrose, glucose and fructose was performed using HPLC with the methodology by Carranza-Saavedra, et al. (2025).

### 2.6. Flavonoid solubility in M9 medium and kinetic parameters

Flavonoid solubility test of naringenin was performed using the methodology by Carranza-Saavedra, et al. (2025). Maximum specific growth rate, substrate yields in biomass, substrate in product, and biomass in product were calculated following the methodology outlined by Carranza-Saavedra et al. (2021).

### 2.7. Statistical analysis

Kinetic parameters and production results obtained from experiments were analyzed against each other using Duncan’s multiple comparison (Duncan, 1955) with a confidence level of 95%. All experiments were performed unless in duplicate and data obtained were expressed as mean ± standard error.

## 3. Results and discussion

### 3.1. Establishing a platform for flavonoid 7-O-glycosylation

The development of an efficient microbial platform for the production of flavonoid glucosides remains a significant industrial challenge. Building upon our previous work with the *E. coli* W-derived strain SBG1869, which carries mutations in the *xyl*A, *pgm*, *zwf*, and *glg*C genes, we established a system that exploits a unique metabolic configuration using sucrose as a carbon source: preferential catabolism of fructose for biomass generation, while simultaneously channeling glucose to increase intracellular UDP-glucose (UDPG) levels, thereby enabling efficient glycosylation reactions (Fig. 2A) (Carranza-Saavedra et al., 2025). In earlier studies, this strategy achieved the highest reported titers of chrysin-7-O-glucoside usin the glycosyltransferase *Yji*C from *Bacillus licheniformis* (Carranza-Saavedra et al., 2025). However, chrysin’s poor solubility may have limited the observable efficiency of the platform, potentially masking its full capacity to glycosylate flavonoids.

**Fig. 2.**
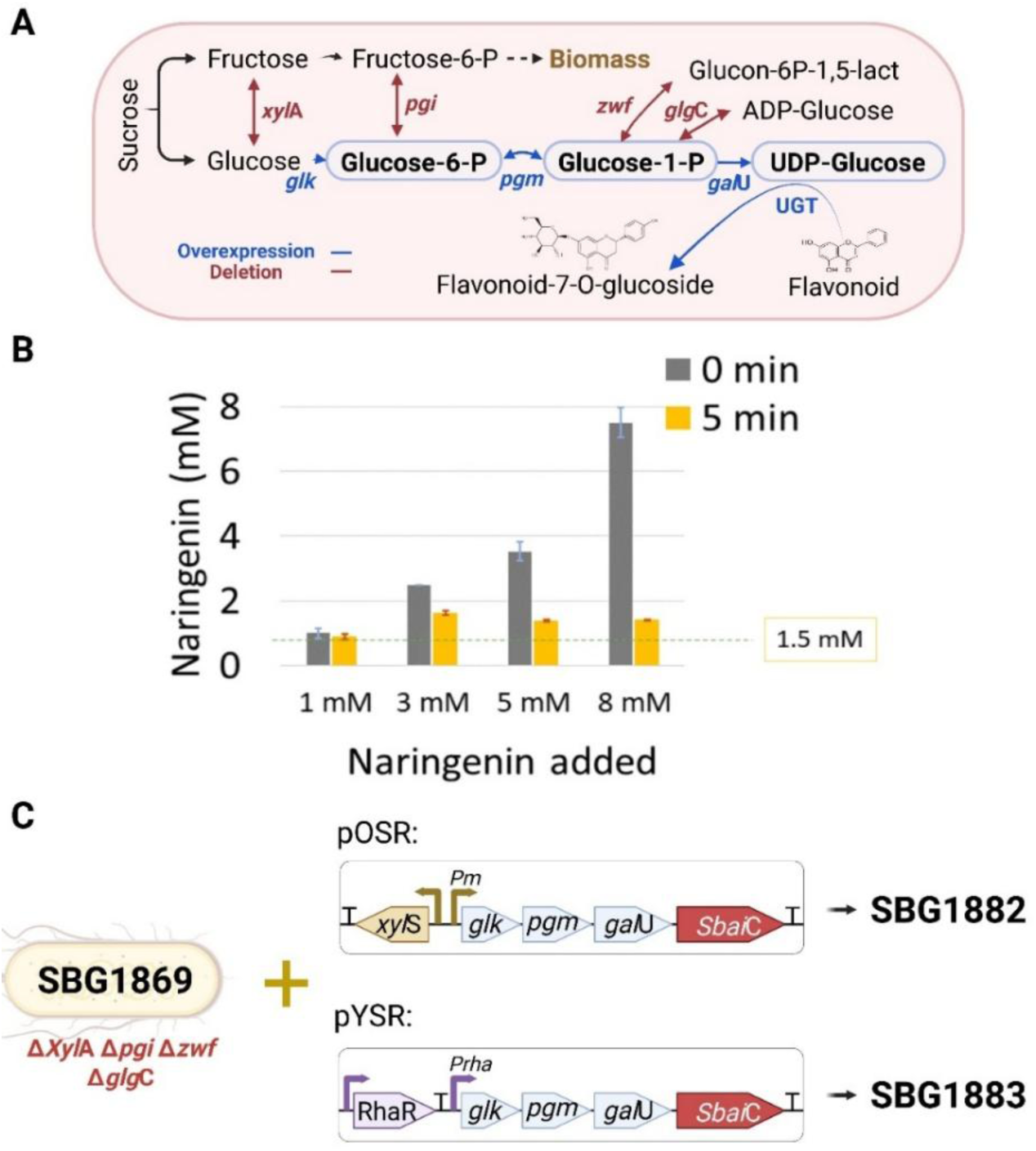
Scheme of engineered *E. coli* W platform for UDPG biosynthesis and flavonoid 7-O-glycosylation. **A.** Metabolic design of the SBG1869 chassis (Δ*xyl*A-*pgi*-*zwf*-*glg*C) highlighting the preferential use of fructose for biomass and excretion of glucose, which is recycled for UDPG biosynthesis via overexpression of *glk*, *pgm* and *gal*U. UDPG is then utilized by the plant-derived glycosyltransferase (UGT) for flavonoid glycosylation. Test of solubility of naringenin in M9 medium with 10 g/L sucrose at 37 °C and without cells. **C.** Genetic constructs used in strains SBG1882 (*XylS*/*Pm* system) and SBG1883 (*RhaR*/*Prha* system), both expressing the UDPG biosynthetic genes and *SbaiC*7OGT under medium–low copy number origin of replication (pBBR).

To more accurately assess the system’s potential, naringenin was selected as the substrate for the production of naringenin-7-O-glucoside (prunin). This choice was based not only on its role as a biologically relevant molecule and core structural scaffold in flavonoid biosynthesis (Rencoret et al., 2022; Tang et al., 2025), making it a versatile precursor for synthesizing a wide range of related flavonoids, but also on its superior solubility (Fig. 2B) and lower cytotoxicity compared to chrysin (Carranza-Saavedra et al., 2025). These attributes made it an ideal candidate to demonstrate the practical potential of our engineered *E. coli* strain SBG1869 (Fig. 2A). It is worth noting that at higher naringenin concentrations, solubility would become a limiting factor for glycosylation (Fig. 2B). Therefore, a concentration of 3 mM naringenin, corresponding to an effective bioavailable concentration of approximately 2 mM in the medium, was selected for prunin production.

To enable prunin synthesis, we expressed the *SbaiC*7OGT gene, a UGT from *S. baicalensis*, in *E. coli* SBG1869. This glycosyltransferase specifically catalyzes the 7-O-glycosylation of flavonoids (Hirotani et al., 2000). The gene was co-expressed with a UDPG biosynthetic cluster (*glk*, *pgm*, and *gal*U) using plasmids regulated by the *XylS*/*Pm* (strain SBG1882) and *RhaR*/*Prha* (strain SBG1883) systems, both maintained under medium-low copy number origins of replication (pBBR; Fig. 2C). Initially, we observed a markedly low transformation efficiency compared to previous studies (Carranza-Saavedra et al., 2025), suggesting potential genetic instability caused by *SbaiC*7OGT-associated toxicity. After overcoming this challenge, we were able to isolate transformants corresponding to strains SBG1882 and SBG1883. Post-bioprocess analysis using PCR with the R24 and PS2 primers, designed to amplify the full transcriptional unit (Fig. 3A), revealed plasmid heterogeneity and the emergence of mixed populations (Fig. 3B, 3C), likely due to metabolic burden or improper protein folding resulting from *SbaiC*7OGT overexpression.

**Fig. 3.**
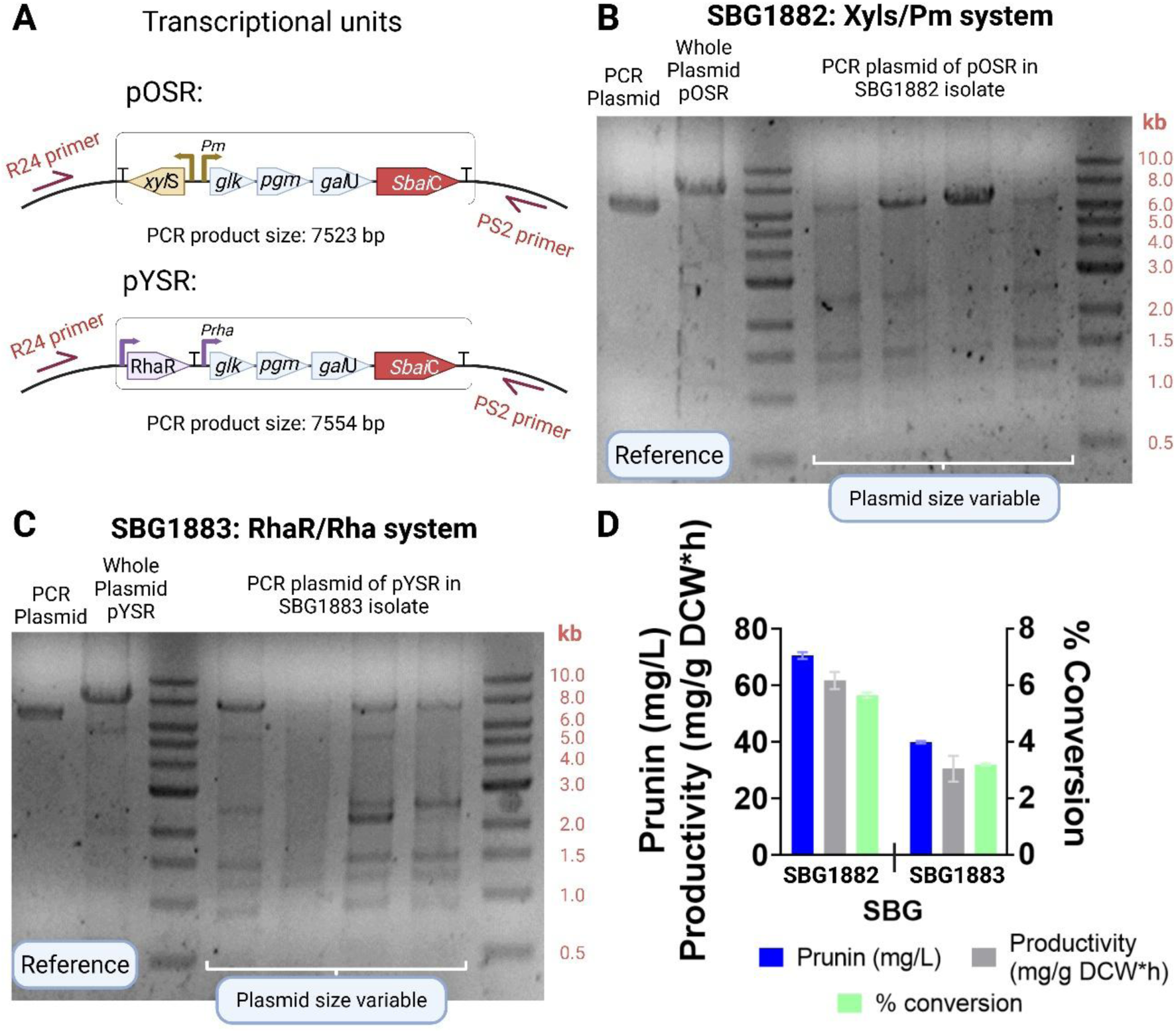
Evaluation of plasmid stability and prunin production using *XylS*/*Pm* (SBG1882) and *RhaR*/*Prha* (SBG1883) expression systems with *SbaiC*7OGT. **A.** Schematic representation of the transcriptional units pOSR (XylS/Pm) and pYSR (RhaR/Prha), each harboring the UDP-glucose biosynthetic genes (glk, pgm, galU) and the glycosyltransferase gene *SbaiC*7OGT under their respective promoters. PCR primers R24 and PS2 were designed to amplify the full transcriptional unit (7.5 kb approx.); **B**-**C.** Gel electrophoresis of whole plasmid preparations and PCR-amplified transcriptional units from isolates of SBG1882 (**B)** and SBG1883 **(C)**. Variable plasmid sizes and band intensities indicate heterogeneity and instability within the engineered strains; **D.** Prunin titers (blue), specific productivity (gray), and naringenin to prunin conversion efficiency (green) achieved in SBG1882 and SBG1883 strains.

Despite these challenges, prunin was produced at concentrations of 39.9 ± 0.4 mg/L and 70.5 ± 1.2 mg/L, with conversion efficiencies of 3.2 ± 0.1% and 5.6 ± 0.3% using the *XylS*/*Pm* and *RhaR*/*Prha* systems, respectively (Fig. 3D). These relatively low conversion rates, especially when compared to other flavonoids (Dorjjugder & Taguchi, 2022; Koirala et al., 2019; H. Li et al., 2022; Thapa et al., 2019), highlight the need for tighter transcriptional control. This is particularly relevant given that chrysin, a poorly soluble flavonoid, still achieved higher conversions, underscoring that the limited prunin yields are more likely due to suboptimal *SbaiC*7OGT expression than substrate bioavailability. Thus, enhanced regulatory systems are therefore essential to stabilize gene expression and improve glycosylation performance.

### 3.2. Mitigating genetic instability and cytotoxicity in *Sbai*C7OGT expression

In industrial-scale fermentation, achieving reproducibility and high productivity requires maintaining a delicate balance between cellular metabolic capacity and synthetic gene expression. Given the limitations of conventional chemically inducible systems, we implemented a novel phosphate-responsive transcriptional system, *Pliar*53, previously developed by our group (Torres-Bacete et al., 2021), while this system has proven effective for self-regulation in two-component regulatory networks (Righetti et al., 2021; Righetti & Kahramanoğulları, 2021), it has not yet been applied to microbial bioprocesses aimed at producing complex biomolecules. In this study, we harnessed the *Pliar*53 system to enable self-inducible gene expression by Pi availability in the culture medium, offering a cost-effective and scalable alternative to traditional induction methods.

To evaluate the performance of this regulatory system, we first constructed the SBG1888 platform, optimized for efficient UDPG biosynthesis. This strain combined the parental chassis SBG1869 with the pUDG plasmid encoding the *glk*, *pgm*, and *gal*U genes (Fig. 4A). For consistent expression analysis and fair comparison with conventional systems, all transcriptional units were standardized with the same ribosome binding site (RBS; STD) and a low-copy origin of replication (RK2). The *SbaiC*7OGT gene was then expressed in SBG1888 under the control of the phosphate-responsive *Pliar*53 promoter, resulting in strain SBG2339 (Fig. 4B). This design aimed to minimize leaky expression of *SbaiC*7OGT, which could otherwise induce cellular toxicity. To ensure repression of gene expression during plasmid construction, strain propagation, and pre-inoculum preparation, all media used with the *Pliar*53 system were supplemented with 5 mM phosphate, concentration found to not allow transcription (Torres-Bacete et al., 2021) (Fig. 4C). Induction of gene expression was triggered only during the production phase, using phosphate-depleted media (M3 medium) supplemented with yeast extract (YE), an industrially relevant additive (see Methods for details).

**Fig. 4.**
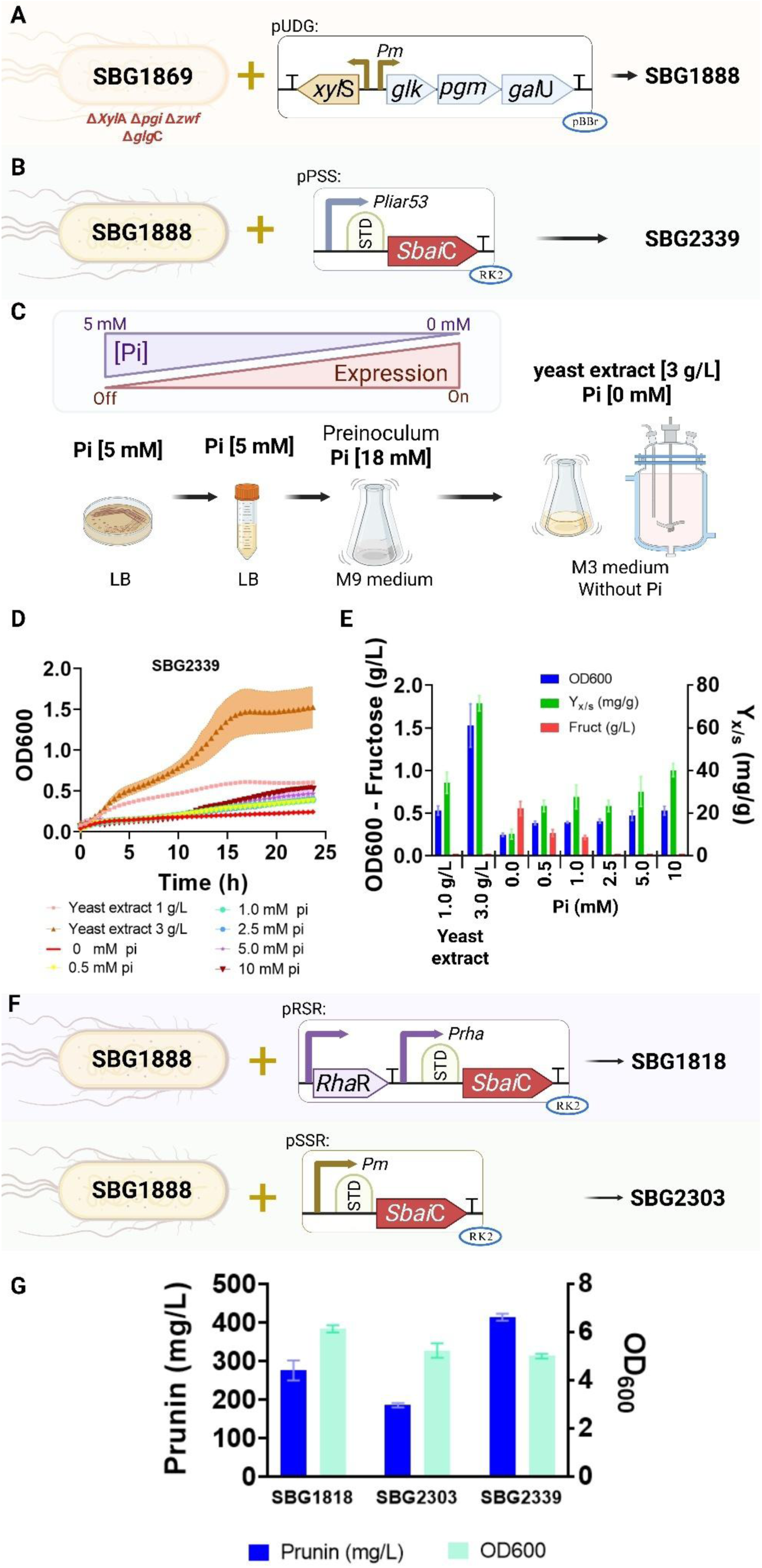
Construction of synthetic *E. coli* W platforms for UDPG biosynthesis and regulated expression of *SbaiC*7OGT. **A.** Engineering of strain SBG1888 by combining the parental strain SBG1869 with plasmid pUDG, which overexpresses the UDPG biosynthetic genes (*glk*, *pgm*, *gal*U) under the control of the *Pm* promoter and *xylS* regulator; **B.** Design of *SbaiC*7OGT expression using STD RBS, phosphate-limitation responsive *Pliar*53 (SBG2339), all integrated into SBG1888 and maintained on low-copy RK2 origin plasmids; **C.** Schematic of the cultivation strategy used to repress and then induce expression of *SbaiC*7OGT by modulating phosphate (Pi) availability. Repression was maintained in LB and pre-inoculum media supplemented with 5-18 mM Pi. Expression was triggered in production medium with 0 mM pi and 3 g/L yeast extract; **D.** Growth profiles of SBG2339 under different Pi concentrations and yeast extract levels; **E.** OD600, fructose levels, and biomass yield (Yx/s) after 24 h of SBG2339. **F.** Design of transcriptional control systems for *SbaiC*7OGT expression using STD RBS: *RhaR*/*Prha* (SBG1818) and *Pm* (SBG2303), all integrated into SBG1888 and maintained on low-copy RK2 origin plasmids. **G.** Comparison of prunin production and growth at 24 h among strains with different promoters.

To determine optimal media conditions, we tested varying concentrations of YE and inorganic phosphate (Pi). We found that 3 g/L of YE, without additional Pi supplementation, supported both cell growth and sucrose consumption, resulting in higher OD_600_ and biomass yield (Yx/s) compared to Pi-supplemented conditions (Fig. 4D and 4E). This effect is likely due to the nutrient-rich composition of YE, which is derived from autolyzed *Saccharomyces cerevisiae* and contains amino acids, peptides, nucleotides (AMP, GMP, CMP), B-complex vitamins, and trace minerals, including small amounts of phosphate (Gao et al., 2024; Taowkrue et al., 2024). Its inclusion appeared to buffer the cells against phosphate starvation while still enabling promoter activation.

In contrast, Pi-depleted cultures without YE experienced prolonged lag phases, lower OD_600_, and fructose excretion, likely due to inadequate phosphate availability for basic metabolic functions (Fig. 4E). Interestingly, cultures supplemented with 10 mM Pi exhibited metabolic behavior similar to those containing 1 g/L YE. However, glycosyltransferase transcription was likely repressed under these conditions, as this Pi concentration exceeds the activation threshold of the *Pliar*53 system (Torres-Bacete et al., 2021).

Regarding prunin production, titers obtained with the phosphate-responsive *Pliar*53 system (strain SBG2339) were compared to those achieved using the conventional *RhaR*/*Prha* (SBG1818) and *XylS*/*Pm* (SBG2303) promoters (Fig. 4F). The *Pliar*53 system delivered the highest prunin titer, reaching 413.9 ± 8.9 mg/L at 24 h, outperforming the *RhaR*/*Prha* and *XylS*/*Pm* systems, which produced 275.3 ± 26.0 mg/L and 185.4 ± 5.8 mg/L, respectively (Fig. 4G). These results underscore the superior performance of the *Pliar*53 expression platform. It is worth noting that fractionating the transcriptional unit into two separate modules *(i)* one for UDPG biosynthesis and *(ii)* another for *SbaiC*7OGT expression, when combined with the *RhaR*/*Prha* or *XylS*/*Pm* systems, yielded higher prunin concentrations than when the entire cluster was expressed as a single unit (Fig. 2C and Fig. 3D). Based on these results, all subsequent experiments employed this dual-module strategy, using *XylS*/*Pm* for UDPG biosynthesis and *Pliar*53 for prunin production.

These findings confirm *Pliar*53 as a robust and efficient system for glycosyltransferase expression under phosphate-limited conditions. Furthermore, its tight transcriptional control opens opportunities for future improvements, such as incorporating stronger RBSs or increasing plasmid copy number, approaches that are often constrained in conventional systems due to metabolic burden or genetic instability.

### 3.3. Enhancing translational efficiency to increase prunin production

Optimizing translation through strong promoters, efficient ribosome binding sites (RBS), and appropriate plasmid copy numbers is critical for maximizing protein expression in heterologous hosts. Inefficient translation can result in low yields, poor resource utilization, and increased metabolic burden, ultimately limiting the production of high-value compounds such as glycosylated flavonoids (Watts et al., 2021). To evaluate the impact of translational elements on *SbaiC*7OGT expression and glycosylation efficiency, we generated variants of the *Pliar*53 expression system by modulating RBS strength and plasmid origin. The previously constructed strain SBG2339, carrying a standard medium-strength RBS (STD), served as the baseline. We then introduced the stronger BCD2 RBS variant (Blázquez et al., 2023) into the same *Pliar*53 system, yielding strain SBG2338 (Fig. 5A).

**Fig. 5.**
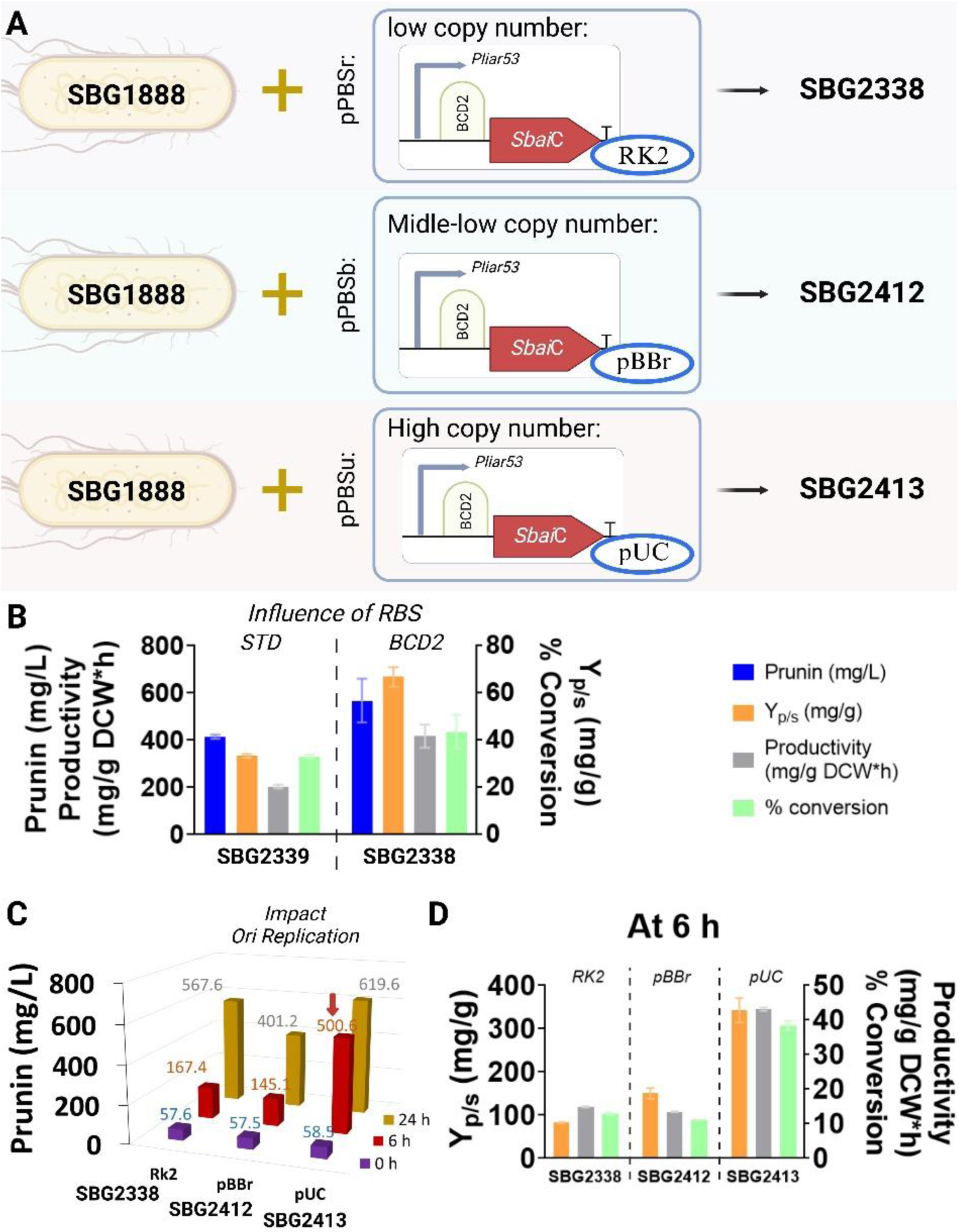
Impact of translational control elements on prunin production using the *Pliar53* system. **A.** Genetic design of SBG1888-derived strains expressing *SbaiC*7OGT with different plasmid copy numbers: low-copy (RK2, SBG2338), medium-copy (pBBR1, SBG2412), and high-copy (pUC, SBG2413); **B.** Comparison of prunin production, productivity, yield (Yp/s) and % conversion among strains with different RBS using the *Pliar*53 system at 24 h; **C.** Prunin production profiles of the three strains with different origin of replications at 0, 6 and 24 h. **D.** Product yield (Yp/s), productivity (mg/g DCW·h), and % conversion of naringenin to prunin at 6 h.

Incorporation of the BCD2 RBS led to a marked improvement in prunin production, with SBG2338 reaching 567.6 ± 9.3 mg/L at 24 h, representing a 37% increase compared to the STD RBS. This enhancement was accompanied by higher naringenin-to-prunin conversion (43%), increased product-to-substrate yield (Yp/s), and greater specific productivity (Fig. 5B). These results highlight the strong potential of the BCD2 RBS, despite its limited use in metabolic engineering due to its tendency to impose metabolic burden (Carranza-Saavedra et al., 2023), as a powerful translational element for boosting heterologous gene expression.

Following the enhancement of translation with the BCD2 RBS, we next evaluated how plasmid copy number affected prunin biosynthesis. Strains were constructed carrying the strong BCD2 RBS combined with either a medium-copy origin (pBBR1) or a high-copy origin (pUC), yielding SBG2412 and SBG2413 respectively (Fig. 5A). Overall sucrose consumption and glucose release were comparable across strains at 24 h (Supplementary Figure S1), but prunin production showed clear differences. The low-copy control strain SBG2338 produced about 567.6 ± 9.3 mg/L, whereas SBG2412 harboring an additional pBBR1 plasmid reached only 401 ± 14.2 mg/L. In contrast, the high-copy pUC strain SBG2413 achieved the best performance with 620 ± 10.0 mg/L corresponding to a 48% conversion rate (Fig. 5C). Early time-point measurements further highlighted the advantage of high copy number. By 6 h SBG2413 had already accumulated over 501 ± 17.7 prunin, threefold higher than the medium-copy strains. Altogether, these results show that coupling BCD2 with a high-copy plasmid markedly improved productivity, yield (Yp/s), and conversion efficiency (Fig. 5D).

Although co-expressing two plasmids with the pBBR1 origin in *E. coli* W proved feasible, it caused a modest growth delay at 24 h. More importantly, despite the known drawbacks of high-copy plasmids like pUC, often linked to metabolic burden and instability (Carranza-Saavedra et al., 2023), the *Pliar*53-based system maintained stable *SbaiC*7OGT expression without growth defects. These results establish SBG2413 as a robust and scalable microbial chassis for high-yield flavonoid glycoside production, combining effective translational tuning with metabolic resilience under optimized bioprocess conditions.

### 3.4. Optimization of the prunin production in fed-batch with SBG2413 in bench-scale bioreactor

The scale-up of bioprocesses is critical for enhancing the production titers of target biomolecules. To enable effective transition to bench-scale bioreactor fermentation, preliminary fed-batch strategies were evaluated in shake flasks to characterize kinetic behavior and guide optimization decisions. To improve prunin production at flask scale, two fed-batch strategies were implemented using the SBG2413 strain in M3 medium, each involving sequential feeding at 6 and 12 h. In the first strategy (Fig. 6A), YE (3 g/L) and naringenin 817 mg/L (3 mM) were added at 6 h, followed by sucrose (10 g/L) at 12 h. In the second (Fig. 6B), phosphate (Pi, 5 mM) and naringenin 817 mg/L (3 mM) were added at 6 h, with sucrose (10 g/L) introduced at 12 h.

**Fig. 6.**
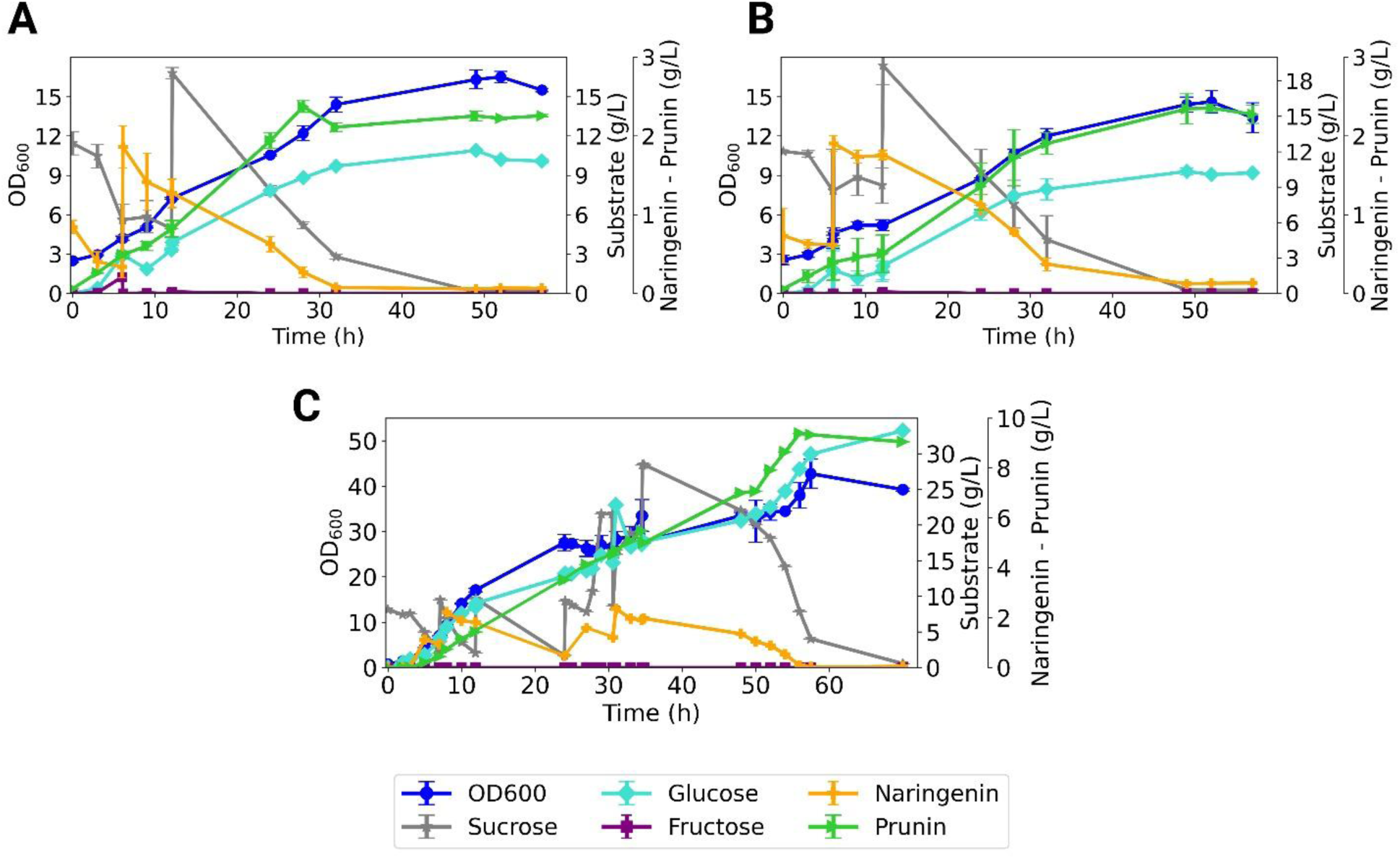
Fed-batch strategies to optimize prunin production in *E. coli* SBG2413. **A.** Shake flask fed-batch with yeast extract and naringenin addition at 6 h and sucrose at 12 h. **B.** Shake flask fed-batch with pi and naringenin at 6 h and sucrose at 12 h. **C.** Fed-batch bioreactor profile of SBG2413 under phosphate-limited conditions with sequential naringenin pulses and sucrose feeding.

The YE-based strategy yielded the highest prunin concentration in flask shake, reaching 2373±74.4 mg/L with a conversion efficiency of 54.6% at 28 h (Fig. 6A). This result indicates that the complex nitrogen sources and growth factors present in yeast extract enhanced both biomass accumulation and enzymatic glycosylation activity. In contrast, the Pi-based strategy achieved a slightly lower titer of 2345±189.8 mg/L and 54.0% conversion, but only at a much later timepoint of 49 h (Fig. 6B). Both approaches enabled effective sucrose consumption and substrate conversion, yet the yeast extract-fed system exhibited a more favorable growth-productivity profile. The earlier and more rapid increase in prunin concentration suggests that *SbaiC*7OGT expression was induced earlier and more efficiently in the presence of YE, highlighting the value of nutrient-rich supplements in improving catalytic performance, particularly in resource-constrained, small-scale setups.

Encouraged by these results, we scaled up the process to a bioreactor using the SBG2413 strain under phosphate-limited, fed-batch conditions with YE (Fig. 6C). Cell growth followed a typical fed-batch profile, reaching an OD600 of 50.4 ± 3.8 by 57 h, which corresponded to approximately 22.7 g CDW/L. Sucrose, serving as the primary carbon source, was rapidly consumed after each feeding pulse, with transient accumulation observed beyond 30 h, coinciding with a reduction of the maximum specific growth rate (µmax) from 0.41 h⁻¹ to 0.12 h⁻¹. This metabolic slowdown suggested carbon overflow and reduced efficiency of sucrose utilization under high cell density.

Naringenin was pulsed at defined intervals and consistently converted into prunin with 70% efficiency, confirming the system’s capacity to sustain catalysis under repeated substrate loading. The process was intentionally stopped at 56 h, as both growth and productivity had plateaued and further substrate addition was no longer efficiently converted. At this point, final prunin concentration reached 9387 mg/L, corresponding to a yield (Yp/s) of 0.42 g/g sucrose and a biomass yield (Yx/s) of 0.21 g/g sucrose. Specific productivity peaked at 151 mg/g DCW·h during the exponential production phase (12–30 h) and gradually declined thereafter as cells entered stationary phase.

Interestingly, between 12 and 24 h might the sucrose supplementation was required. Although, as the initial feed was sufficient to sustain growth and product formation, this interval represents a critical window where biomass accumulation and early induction of *SbaiC*7OGT occur in parallel, suggesting that future optimization could involve fine-tuning feeding schedules or continuous-feeding strategies to balance growth and glycosylation rates more effectively.

Importantly, the use of a high-copy pUC-origin plasmid in SBG2413 did not impose detectable metabolic burden or growth defects throughout the 60-h run. This stability emphasizes the advantage of the *Pliar*53 system in tightly regulating expression of potentially toxic genes under fed-batch conditions. Overall, the combination of phosphate-responsive control, sucrose-based feeding, and translationally optimized SBG2413 enabled the highest reported prunin titer (>9 g/L) with efficient conversion in bench-scale bioreactors. These results validate the scalability of the platform and lay the foundation for further optimization through dynamic feeding strategies, continuous processing, or scale-up to industrial fermenters.

### 3.5. Broadening platform versatility: proof of concept with diverse flavonoids in shake flasks

The production of structurally diverse flavonoid glycosides is highly valuable for pharmaceutical, cosmetic and food applications. Variations in backbone structure, hydroxylation patterns and physicochemical properties influence bioactivity, bioavailability and microbial tolerance, making the development of robust production platforms a key challenge in industrial biotechnology.

To assess the versatility of our engineered *E. coli* W platform (strain SBG2413, derived from SBG1869) we evaluated its capacity to glycosylate five structurally distinct flavonoids including daidzein (isoflavone), pinocembrin (flavanone), hesperetin (flavanone), apigenin (flavone) and chrysin (flavone). All reactions were performed in shake flasks. As shown in Table 2, four of the five compounds were converted to their corresponding 7-O-glucosides with conversion efficiencies exceeding 40 percent. These included pinocembrin-7-O-glucoside, apigenin-7-O-glucoside (apigetrin), hesperetin-7-O-glucoside and chrysin-7-O-glucoside. Titers at 24 h ranged from 608.2 ± 26.7 mg/L for hesperetin to 795.8 ± 104.9 mg/L for chrysin. In contrast, daidzein-7-O-glucoside reached only 263.6 ± 5.1 mg/L, likely due to its poor solubility and previously reported cytotoxic effects in *E. coli* W (Carranza-Saavedra et al., 2025).

**Table 2.**
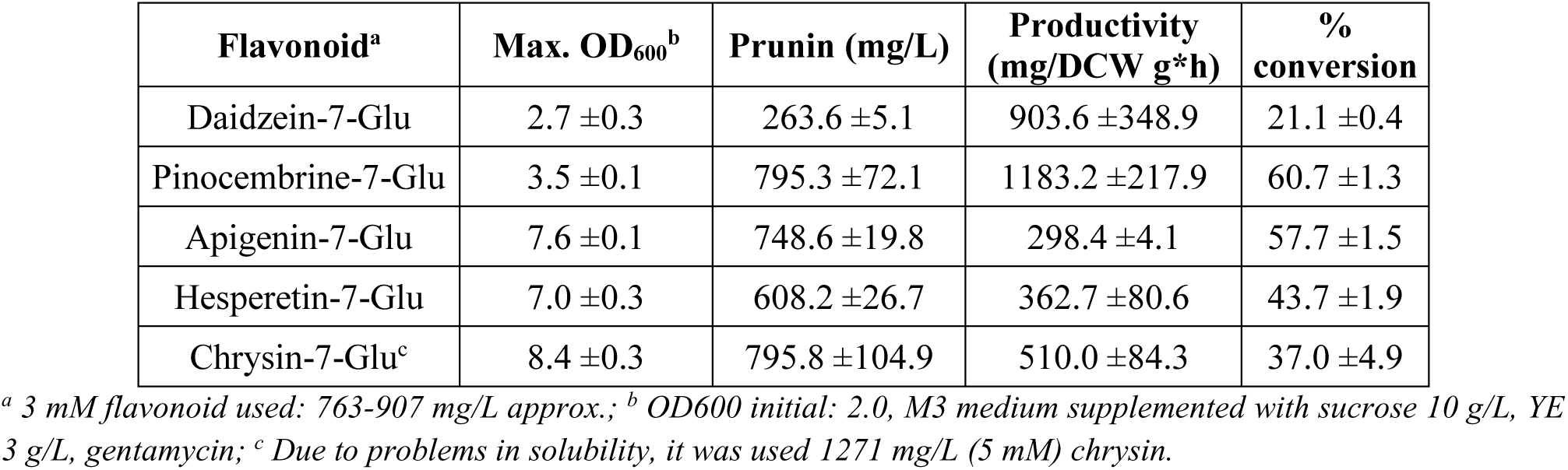
Glycosylation of structurally diverse flavonoids in *E. coli* SBG2413 using *SbaiC*7OGT and *Pliar*1.53 system.

| Flavonoid<sup>a</sup> | Max. OD<sub>600</sub><sup>b</sup> | Prunin (mg/L) | Productivity (mg/DCW g*h) | % conversion |
| --- | --- | --- | --- | --- |
| Daidzein-7-Glu | $2.7 \pm 0.3$ | $263.6 \pm 5.1$ | $903.6 \pm 348.9$ | $21.1 \pm 0.4$ |
| Pinocembrine-7-Glu | $3.5 \pm 0.1$ | $795.3 \pm 72.1$ | $1183.2 \pm 217.9$ | $60.7 \pm 1.3$ |
| Apigenin-7-Glu | $7.6 \pm 0.1$ | $748.6 \pm 19.8$ | $298.4 \pm 4.1$ | $57.7 \pm 1.5$ |
| Hesperetin-7-Glu | $7.0 \pm 0.3$ | $608.2 \pm 26.7$ | $362.7 \pm 80.6$ | $43.7 \pm 1.9$ |
| Chrysin-7-Glu<sup>c</sup> | $8.4 \pm 0.3$ | $795.8 \pm 104.9$ | $510.0 \pm 84.3$ | $37.0 \pm 4.9$ |

<sup>a</sup> 3 mM flavonoid used: 763-907 mg/L approx.; <sup>b</sup> OD<sub>600</sub> initial: 2.0, M3 medium supplemented with sucrose 10 g/L, YE 3 g/L, gentamycin; <sup>c</sup> Due to problems in solubility, it was used 1271 mg/L (5 mM) chrysin.

Biomass formation varied across substrates. The highest OD₆₀₀ values were obtained with chrysin (8.4 ± 0.3) and apigenin (7.6 ± 0.1), suggesting better cellular tolerance despite chrysin’s relatively lower conversion. Daidzein cultures exhibited the lowest OD₆₀₀ (2.7 ± 0.3), consistent with its inhibitory effects on growth.

Productivity patterns largely mirrored conversion trends. Pinocembrin-7-O-glucoside achieved the highest specific productivity (1183.2 ± 217.9 mg/g DCW·h), followed by chrysin-7-O-glucoside (510.0 ± 84.3 mg/g DCW·h). Daidzein’s low productivity (903.6 ± 348.9 mg/g DCW·h) reflected the combined impact of low cell density and poor substrate solubility.

Overall, correlations between OD₆₀₀, productivity and conversion efficiency indicate that both substrate toxicity and physicochemical properties strongly shape glycosylation performance. These results validate the broad substrate compatibility of the SBG2413 platform and the catalytic versatility of *SbaiC*7OGT, highlighting its robustness under varying substrate conditions and supporting its potential for expansion into additional flavonoid subclasses and industrial-scale bio-based synthesis of high-value glycosides.

### 3.8. Discussion

The microbial production of flavonoid glycosides has attracted growing attention due to their improved physicochemical properties, enhanced bioactivity, and potential applications in pharmaceuticals, nutraceuticals, and functional foods. However, the efficient biosynthesis of 7-O-glucosides, has remained challenging because of the low solubility and toxicity of the aglycone precursors, limited enzyme activity, and insufficient metabolic robustness of existing microbial platforms. In this study, we established a phosphate-responsive system that allows dynamic control of a toxic UDP-glycosyltransferase and enables the efficient biosynthesis of prunin at titers exceeding 9 g/L, a production level that surpasses previous microbial reports by more than two orders of magnitude (Dorjjugder & Taguchi, 2022; Koirala et al., 2019; H. Li et al., 2022; Thapa et al., 2019).

Earlier attempts to produce flavonoid 7-O-glucosides relied on heterologous expression of plant glycosyltransferases in *E. coli*, but these systems typically suffered from modest yields and limited scalability. For instance, it was reported the conversion of several flavonoids into their 7-O-glucosides using a tobacco-derived enzyme expressed in *E. coli*, with final titers ranging from 24 to 118 mg/L, depending on the substrate, and requiring sequential substrate feeding to avoid precipitation (Dorjjugder & Taguchi, 2022). Similarly, biotransformation approaches employing probiotic strains such as *Bacillus amyloliquefaciens* resulted in the formation of prunin and other derivatives (Koirala et al., 2019), but production remained at the milligram-per-liter scale and lacked the robustness needed for industrial implementation. In contrast, our platform achieved gram-per-liter titers directly from sucrose in controlled bioreactors, highlighting the advantage of integrating a novel expression system enabling scalable production like the *Pliar*53 with efficient translation a good metabolic burden balance.

Recent strategies have also explored the use of glycosylation not as a final modification but as a metabolic engineering tool to alleviate solubility and toxicity issues during flavonoid biosynthesis. Engineered *Saccharomyces cerevisiae* to transiently channel naringenin flux into prunin, thereby improving secretion and cell growth. After extracellular hydrolysis, the strain reached 1184 mg/L of naringenin, representing a 7.9-fold increase compared with the parental strain (H. Li et al., 2022). This elegant approach underscores the value of glycosylation as a physiological strategy. Nevertheless, while glycosylation in that case was an auxiliary step toward aglycone accumulation, in our system the glycosylated form itself is the target product, thereby providing direct access to bioactive molecules with superior stability and solubility.

Enzymatic cascade systems have also been developed to overcome the cost and supply limitations of sugar donors for glycosylation. For example, it was established an *in vitro* UDP-recycling system based on sucrose synthase coupled with glycosyltransferases, enabling the synthesis of multiple naringenin glucosides and quercetin rhamnosides with high conversion efficiencies (Thapa et al., 2019). While such approaches elegantly demonstrate the potential of enzymatic catalysis, they remain dependent on expensive cofactors and purified proteins, limiting their scalability for industrial fermentation. By contrast, our microbial platform embeds the entire glycosylation machinery within a robust cellular environment, obviating the need for external cofactors or sequential substrate additions.

Beyond its unprecedented productivity, the *Pliar*53 system consistently outperformed conventional regulators in terms of titer, yield, and productivity, underscoring its suitability for industrial-scale processes where chemical inducers are either cost-prohibitive or toxic. A critical feature of this system was the ability to fine-tune phosphate concentrations in the medium through yeast extract supplementation. Yeast extract not only buffered phosphate release to control gene expression but also simultaneously supported biomass accumulation, thereby fulfilling a dual role as inducer modulator and growth supplement. Remarkably, a single additional dose of yeast extract (3 g/L, reaching a total of 6 g/L) was sufficient to maximize production, a level considerably lower than nutrient-rich conditions used in prior flavonoid glycosylation studies, yet yielding superior titers, yields, and productivities (Liu et al., 2023; Wu et al., 2022). This observation highlights how carefully calibrated nutrient supplementation can replace more complex induction strategies while reducing process costs. In addition, the combination of yeast extract with naringenin further improved platform functionality by addressing a key bottleneck in flavonoid biosynthesis, namely precursor solubility. Compared with chrysin, which exhibited higher cytotoxicity and poorer aqueous compatibility in earlier work (Carranza-Saavedra, 2025), naringenin provided a more soluble and less toxic substrate, which improved reaction kinetics and allowed a more accurate assessment of system potential. This result has broader implications since the physicochemical properties of flavonoid precursors themselves can significantly influence overall system performance and should be considered in the design of future glycosylation workflows.

At the translational level, we also showed that system performance could be further enhanced through rational expression tuning. Incorporation of the non-conventional BCD2 ribosome binding site substantially boosted prunin titers compared with a standard RBS (Blázquez et al., 2023), demonstrating the value of translational fine-tuning for managing expression of toxic genes. When combined with plasmid copy number modulation, the system tolerated even high-copy vectors such as pUC without imposing metabolic burden, a result rarely achieved in toxic-gene contexts. Although a temporary growth lag was observed when dual pBBR1-based plasmids were co-maintained, sucrose metabolism and product formation remained largely unaffected, indicating that *Pliar*53 regulation can balance plasmid load with metabolic flux. Together, these results reveal a previously unattainable level of flexibility in system design, where induction strength, translational efficiency, and plasmid dosage can be co-optimized without compromising host fitness.

Taken together, our results position the phosphate-regulated expression system as a broadly applicable strategy for the production of glycosylated flavonoids. Beyond prunin, the system enabled the biosynthesis of additional 7-O-glucosides, demonstrating versatility and expanding the chemical space accessible through microbial fermentation. The titers achieved here represent the highest reported to date for prunin and stand in sharp contrast with earlier microbial or enzymatic systems. More importantly, the dynamic control of enzyme expression mitigates the toxicity barrier that has historically limited glycosyltransferase deployment in high-density cultures. These advances highlight the potential of our approach not only for flavonoid glycosides but also for other classes of natural products where solubility and toxicity impede efficient biosynthesis. Future studies may explore extending this strategy to consortia-based systems, integrating further pathway optimizations, and evaluating downstream purification and formulation routes, thereby accelerating the translation of microbial glycoside production into scalable biotechnological applications.

## 4. Conclusion

In this work, we demonstrated a phosphate-responsive regulatory system that enables dynamic control of toxic glycosyltransferases, resulting in unprecedented titers of prunin above 9 g/L. Compared with previous microbial or enzymatic strategies, which typically achieved milligram-per-liter levels, our approach represents a breakthrough in both yield and scalability. Beyond prunin, the system proved versatile for the biosynthesis of other 7-O-glucosides, underscoring its potential as a broadly applicable platform for flavonoid glycosylation. By combining high productivity with robust host performance, this strategy provides a powerful blueprint for the microbial manufacturing of value-added natural products with improved solubility, stability, and bioactivity.

## Author contributions

DCS, Conceptualization, Methodology, Investigation, Data curation, Formal analysis, Writing original draft, JTB. Investigation, writing review and editing. JN, Conceptualization, Investigation, Project administration, Formal analysis, Supervision and Writing review and editing, Funding acquisition.

## Acknowledgements

The authors gratefully acknowledge Jarosław Popłoński and his research group at the Department of Food Chemistry and Biocatalysis, Wrocław University of Environmental and Life Sciences (Poland), for kindly providing the *SbaiC*7OGT gene as well as for their valuable support input throughout this work.

