## Supplementary_1 for "A novel expression system enabling scalable production of glycosylated flavonoids in *Escherichia coli* W using a plant-derived toxic gene"

* Correspondence:

Corresponding Author:

E-mail address:

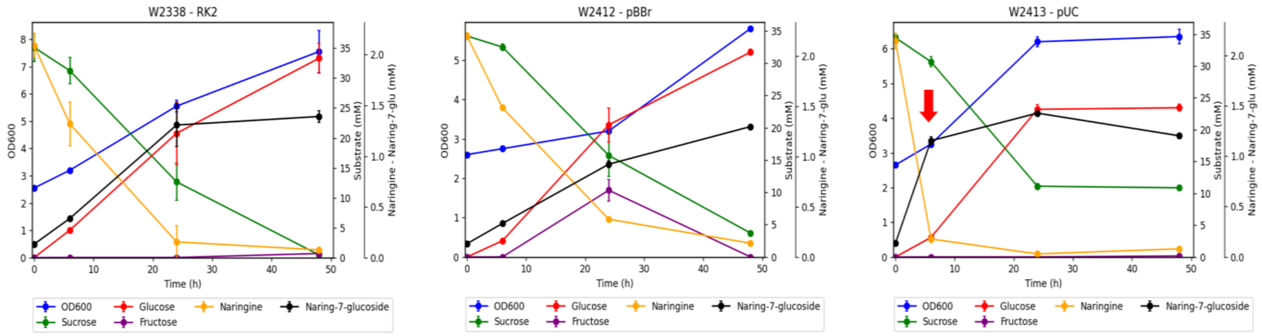


**Supplementary Figure S1.** Kinetics to prunin production in *E. coli* SBG2338 (A), SBG2412 (B) and SBG2413 (C).
